# Mechanism of action of Buqing Granule against Diabetic retinopathy based on network pharmacology and animal experiments

**DOI:** 10.1101/2023.12.20.572649

**Authors:** Yifan Yang, Ling Yuan, Xiangyang Li, Qian Liu, Wenjie Jiang, Taiqiang Jiao, Jiaqing Li, Mengyi Ye, Yang Niu, Yi Nan

**Affiliations:** Ningxia Medical University Key Laboratory of Ningxia Minority Medicine Modernization Ministry of Education, Ningxia Medical University, Yinchuan, Ningxia, China; School of Pharmacy, Ningxia Medical University, Yinchuan, Ningxia Hui Autonomous Region, China; School of Traditional Chinese Medicine, Ningxia Medical University, Yinchuan, Ningxia, China; School of Clinical Medicine, Ningxia Medical University, Yinchuan, Ningxia, China

**Keywords:** Buqing Granule, Diabetic retinopathy, Network Pharmacology, Molecular Mechanisms

## Abstract

**Objective:** For this study, network pharmacology and animal experiments were used together to get a better idea of how BQKL works at the molecular level to treat DR.

**Methods:** In this study, we obtained the relevant action target information of BQKL from the TCMSP and also combined the GeneCards, OMIM, TTD, and PharmGKB databases with the GEO database to obtain the relevant target information of DR. The intersection of these targets was determined using a Venn diagram to identify the target of action for the treatment of DR with BQKL. The target proteins of BQKL for DR were then uploaded to the String database. The resultant data were imported into Cytoscape 3.9.0 to construct PPI networks and identify key targets of action. The DAVID database was used to do a GO and KEGG pathway enrichment analysis of target genes for treating DR with BQKL. Molecular docking was performed to validate the core action targets with the core compounds of BQKL. In addition, we induced DR production in rats by a high-fat, high-sugar diet and intraperitoneal injection of STZ and validated the results obtained from the network pharmacological analysis by changes in body weight and blood glucose, serum levels of biochemical markers, HE staining, immunohistochemistry, qRT-PCR, and Western blot experiments in rats.

**Results:** In this study, quercetin, kaempferol, β-sitosterol, lignanserin, and stigmasterol were identified as the key components, TP53, AKT1, JUN, CASPASE3, MAPK3, and MAPK1 as the key targets, and PI3K-Akt, AGE-RAGE, and MAPK signaling pathways as the main pathways involved. The results of animal experiments showed that BQKL could not only effectively reduce the degree of blood glucose, blood lipids, and oxidative damage in diabetic rats but also slow down the development process of DR. At the same time, it can significantly up-regulate the expression of AKT1, MAPK1, and MAPK3 and down-regulate the expression of CASPASE3, c-JUN, and TP53 in retinal tissue.

**Conclusion:** BQKL ameliorates oxidative stress, apoptosis, and inflammation due to hyperglycemia-related stress by regulating key targets of CASPASE3, AKT1, c-JUN, TP53, MAPK1, and MAPK3, thereby delaying the onset and progression of DR.

## 1. Introduction

Diabetic retinopathy (DR) is a disease that causes damage to the retina due to the deleterious effects of a prolonged state of hyperglycemia on the blood vessels and tissues of the eye, and it is one of the serious complications that require a high level of attention for diabetics [1]. The pathogenesis of DR is complex, and treatment can be challenging. Currently, the best option for reducing the risk of retinopathy onset and progression is to control blood glucose, blood pressure, and lipids [2]. The most common clinical treatments include steroid medications, anti-vascular endothelial growth factor therapy, vitrectomy, and laser therapy [3]. However, these modalities of treatment can be costly for patients and may have certain side effects. As a result, there is a growing interest in finding treatment modalities that have low adverse effects, are safe, and effective in DR research.

Buqing Granule (BQKL) is a traditional prescription with a long history that was first mentioned in the Yangshi Jiacang Fang [4]. It is known for its ability to tonify the kidney, benefit the essence, nourish the liver, and brighten the eyes. Our research group has conducted extensive studies on the treatment of ophthalmopathy using various methods such as clinical observation, modern medicinal chemical extraction, pharmacological experiments, animal experiments, and drug-containing serum interventions in cell damage model experiments [5-9]. The research findings indicate that BQKL exhibits strong antioxidant and hypoglycemic effects. It effectively reduces body weight in db/db mice, is bright-eyed, and has demonstrated high safety. However, current research on the pharmacological effects and molecular mechanisms involved in DR treatment with BQKL still needs to be developed.

Network pharmacology is a new field that combines computer science and systems biology to study the mechanism of drug action and the molecular design of multi-target drugs [10]. It focuses on the study of “drug-disease-target gene” interaction networks at a systemic and comprehensive level. Chinese medicines and their formulas are characterised by multi-component, multi-pathway, and multi-target synergistic effects, which have their own characteristics and advantages in the prevention and treatment of complex diseases. There are many similarities between network pharmacology and the holistic concepts of traditional Chinese medicine, which provide a brand-new perspective and solution for the study of the mechanism of action and the material basis of the efficacy of the Chinese medicines’ compound formulas [11]. The aim of this study is to explore the potential mechanism by which BQKL treats DR using a combination of network pharmacology and animal experiments, with the ultimate goal of establishing a foundation for its future clinical application (Fig 1).

**Fig 1.**
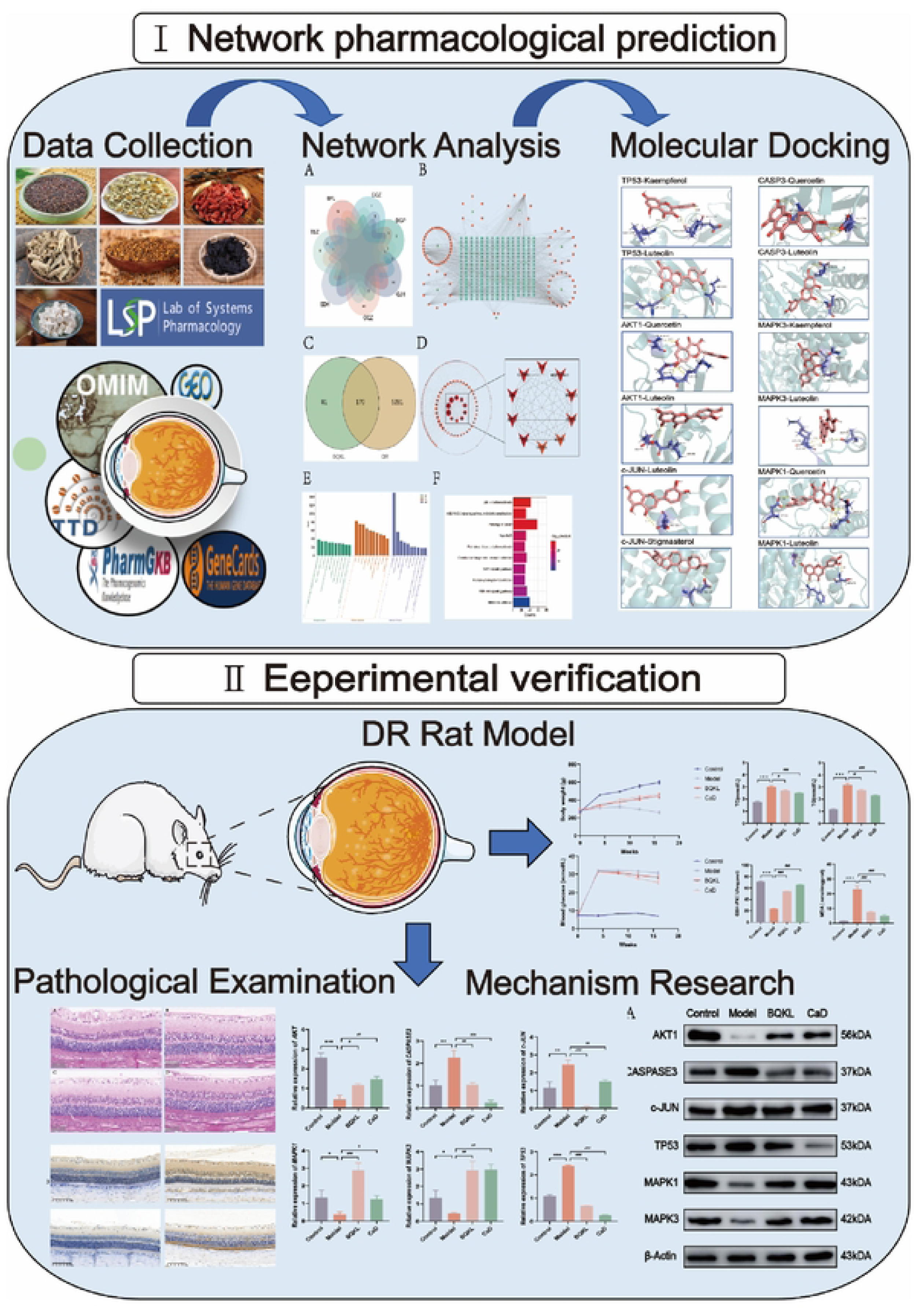
Research Flow Chart.

## 2. Materials and Methods

### 2.1. Screening of the active ingredients in BQKL

The active ingredients in the BQKL were searched using the TCMSP (http://ibts.hkbu.edu.hk/LSP/tcmsp.php) [12] to screen the active ingredients and potential targets of action of the BQKL, which were supplemented by reviewing the literature through PubMed (https://pubmed.ncbi.nlm.nih.gov/) to supplement the literature. A network diagram of “BQKL-Active Ingredients-Targets of Action” was constructed.

### 2.2. Prediction of DR-related targets

Disease targets were collected through GeneCards [13] (https://www.genecards.org/), OMIM [14] (https://omim.org/), TTD [15] (http://db.idrblab.net/ttd/), PharmGKB [16] (https://www.pharmgkb.org/) combined with GEO [17] (http://www.ncbi.nlm.nih.gov/geo/) and other databases to collect disease targets, merge disease targets and de-emphasize them to identify potential targets of action in DR.

### 2.3. Construction of Venn diagrams and PPI networks

InteractiVenn [18] (http://www.interactivenn.net/index2.html) mapped the disease targets derived from the screening to the targets of the BQKL compounds to take the intersection, and the intersecting targets were considered to be the targets of action of the BQKL for the treatment of DR. The intersected targets were imported into the STRING database [19] (https://cn.string-db.org/) to obtain the protein interaction information. Cytoscape 3.9.0 software was used for visualisation, removing free targets, constructing the PPI network, performing network topology analysis, and calculating the degree value of the targets in the PPI network.

### 2.4. Construction of “BQKL ⁃ active ingredient ⁃ target” network diagrams

The network diagram of “BQKL ⁃ active ingredient ⁃ target” was drawn by the plug-in of Cytoscape 3.9.0 software and visualised and analysed.

### 2.5. GO and KEGG enrichment analyses

The intersection targets that overlapped were added to the DAVID database [20] (https://david.ncifcrf.gov/tools.jsp), and then Gene Ontology (GO) analysis and Kyoto Encyclopaedia of Genes and Genomes (KEGG) enrichment analysis were done on them. The results will be visualised using microbiota.

### 2.6. Molecular docking

The SDF structure files of the core compounds in BQKL were downloaded through the Pubchem (https://pubchem.ncbi.nlm.nih.gov/) database [21]. The high-resolution PDB format files of the core genes of the BQKL compounds were obtained from the PDB database (https://www.rcsb.org) [22] and processed by Pymol software for dehydrogenation and hydrogenation after deletion of the proto-ligands contained in the large-molecule receptors. The processed large and small molecules were imported into AutoDockTools 1.5.7 for molecular docking verification, and the docking results were visualized by Pymol software.

### 2.7. Animals

Eighty-five healthy 5-week-old specific-pathogen-free (SPF) male Sprague Dawley (SD) rats, body mass (220±20) g, were purchased from the Animal Experiment Centre of Ningxia Medical University, Animal Licence No. SYXK (Ning) 2020-0001. During the experimental period, an adequate supply of food and water was ensured. All rats were adaptively fed for one week, and their status was observed daily. The Laboratory Animal Welfare Ethics Committee of the Laboratory Animal Centre of Ningxia Medical University (Ethics No. IACUC-NYLAC-2022-113) approved all animal experiments.

### 2.8. Drug preparation

All the herbs used in this study were purchased from the Traditional Chinese Medicine Hospital of Ningxia Medical University and prepared according to the standard. The granules were composed of *Cuscuta chinensis* Lam. (Tusizi) 20g, *Lycium barbarum* L. (Gouqizi) 20g, *Lycium chinense* Mill. (Digupi) 10g, *Plantago asiatica* L. (Cheqianzi) 10g, *Chrysanthemum morifolium* Ramat. (Ganjuhua) 10g, *Rehmannia glutinosa* Libosch. (Shudihuang) 20g, and *Poria cocos* (Schw.) Wolf (Baifuling) 20g. Yuan Ling of the School of Pharmacy at Ningxia Medical University. The positive drug calcium dobesilate (CaD) is produced by Ningxia Kangya Pharmaceutical Co.

### 2.9. Establishment of a rat model of DR

After a week of acclimatized feeding, 30 rats were randomly selected from 40 rats to be the model group and 10 rats to be the blank group. The blank group was given normal chow, and the model group rats were given high-fat and high-sugar chow [23] (Boaigang-1135DM, Beijing Boai Port Biotechnology Co., Ltd., China). After 4 weeks, all the rats were fasted for 12 hours. The rats in the model group were given a single intraperitoneal injection of 60 mg/kg of streptozotocin (STZ, Boaigang-B2001, Beijing Boai Port Biotechnology Co. Ltd., China) dissolved in sodium citrate buffer. The rats in the blank group were given an intraperitoneal injection of the same amount of sodium citrate buffer. After 72 hours, blood from the tail veins of 30 rats was used to test their random blood glucose levels. Three out of the four random blood glucose levels were ≥ 16.7 mmol/L, which showed that the diabetics model worked [24]. After successful modelling, the rats were randomly divided into the model group, the BQKL group (800 mg/kg/d) and the CaD group (104 mg/kg/d) [25], with 10 rats in each group. For 12 weeks, the rats in the BQKL and CaD groups received medication by gavage, and the rats in the normal group and the model group received an equal amount of physiological saline.

### 2.10. Measurement of body weight and blood glucose

During the experimental period, the rats’ eyeballs, fur color, diet, water intake, urine output, body shape, mental status, and activity were observed and recorded weekly, and the body weights and blood glucose of the rats in each group were measured at the end of the acclimatization feeding for 1 week at 0, 4, 8, 12, and 16 weeks, respectively, and the blood sugar values of the rats in each group were determined by the tail-vein blood sampling method through a glucose meter.

### 2.11. Detection of biochemical indicators

We got the serum from each group of rats and used the ELISA assay to measure its absorbance. We then figured out how much total cholesterol (TC), triglyceride (TG), superoxide dismutase (SOD), malondialdehyde (MDA), and glutathione peroxidase (GPx) were in each sample.

### 2.12. Hematoxylin-Eosin (HE) Staining

All rats were taken by intraperitoneal injection of sodium pentobarbital (65mg/kg) for execution [26]. The rat eyeballs were removed and fixed in a paraformaldehyde solution for paraffin embedding. The wax blocks were cut into 5-μm-thick tissue sections using a sectioning machine, routinely deparaffinized, HE-stained, and photographed under a light microscope [27]. Sections were selected, and the retinal thickness and the number of retinal ganglion cells (RGC) were counted.

### 2.13. Immunohistochemical detection of retinal tissue CASPASE3, JUN, MAPK1, and MAPK3 protein expression

The paraffin-embedded eyeballs will be serially sectioned using a sectioning machine to a thickness of 5μm. Sections were routinely dewaxed for aqueous, antigenic repair. The sections were incubated overnight with primary antibodies and then incubated with secondary antibodies for 2 hours at room temperature for DAB color development. After color development, hematoxylin was restained, and the sections were sealed with neutral gum [28]. Image Pro Plus 6.0 software was used for semi-quantitative analysis of tan deep staining.

### 2.14. qRT-PCR

Retinal tissues were taken from each group, and total retinal RNA was extracted by the Trizol one-step method. After reverse transcription into cDNA samples by reverse transcription reagent, Real-time PCR reaction was performed. The relative expression of each target gene was calculated using GAPDH expression as an internal reference control. The CT values were calculated using the 2^-ΔΔCt^ method [29]. Primers were synthesized by Sangon Biotech (Shanghai) Co. (Table 1).

**Table 1.**
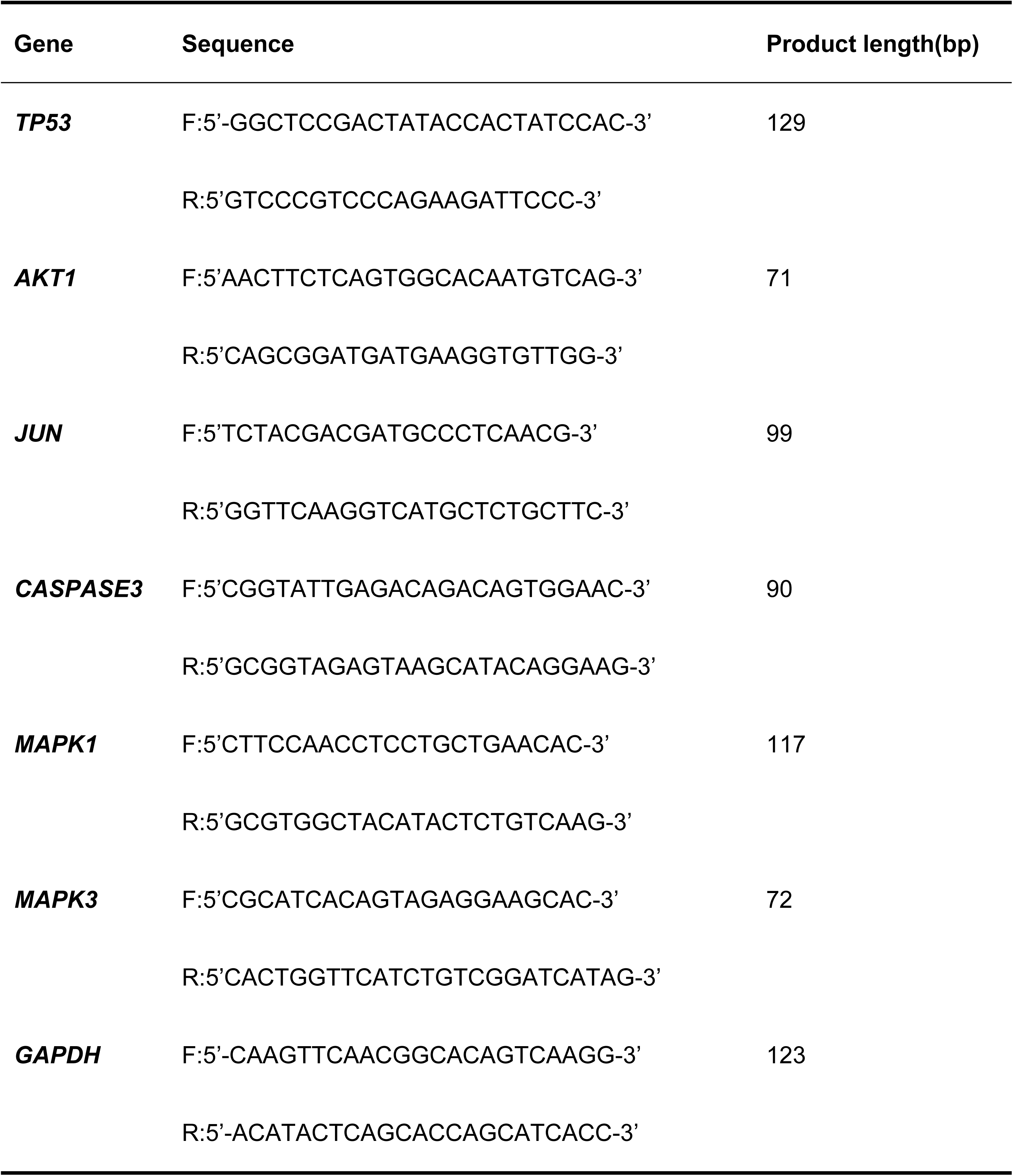
Primer information.

### 2.15. Western blot detection of CASPASE3, AKT1, JUN, TP53, MAPK1, and MAPK3 expression in rat retinal tissues

The retinal tissue was lysed in lysis solution, homogenized well by a homogenizer, and the supernatant was taken after centrifugation for 10 min. The BCA kit method was used to determine the protein content. The samples were separated by SDS-PAGE electrophoresis and transferred to the membrane, which was closed for 2 hours. The primary antibody was added and incubated overnight. The membrane was washed three times and incubated with the corresponding secondary antibody for 2 hours. The color was developed by adding luminescent solution, and the image data were analyzed and processed by Image Pro Plus 6.0 software.

#### Antibodies

MAPK3 Antibody (AF0562, Affinity, US); AKT1 Antibody (AF0836, Affinity, US); c-Jun Antibody (AF6090, Affinity, US); Caspase 3 Antibody (AF6311, Affinity, US); ERK2 Antibody (DF6032, Affinity, US); TP53 Antibody (DF7238, Affinity, US); β-actin (AF7018, Affinity, US); Goat anti-rabbit (S0001, Affinity, US).

### 2.16. Statistical analysis

Data were expressed using the mean ± standard deviation. Comparisons between groups were performed using one-way analysis of variance (ANOVA) and a t-test using GraphPad Prism 9.5.0.

## 3. Results

### 3.1. Results of active ingredient screening and target prediction of BQKL

A total of 11 chemical constituents of *Cuscuta chinensis* Lam., 2 of *Rehmannia glutinosa* Libosch., 45 of *Lycium barbarum* L., 9 of *Plantago asiatica* L., 13 of *Lycium chinense* Mill., 15 of *Poria cocos* (Schw.) Wolf, and 20 of *Chrysanthemum morifolium* Ramat. were collected after screening by searching the literature and consulting the database (Fig 2A). The removal of duplicates was the main active component of BQKL, which totaled 97. After combining and getting rid of duplicate target genes linked to the acquired BQKL compounds, a total of 251 drug targets were found. Cytoscape 3.9.0 software was used to construct the “BQKL-active ingredient-target” (Fig 2B).

**Fig 2.**
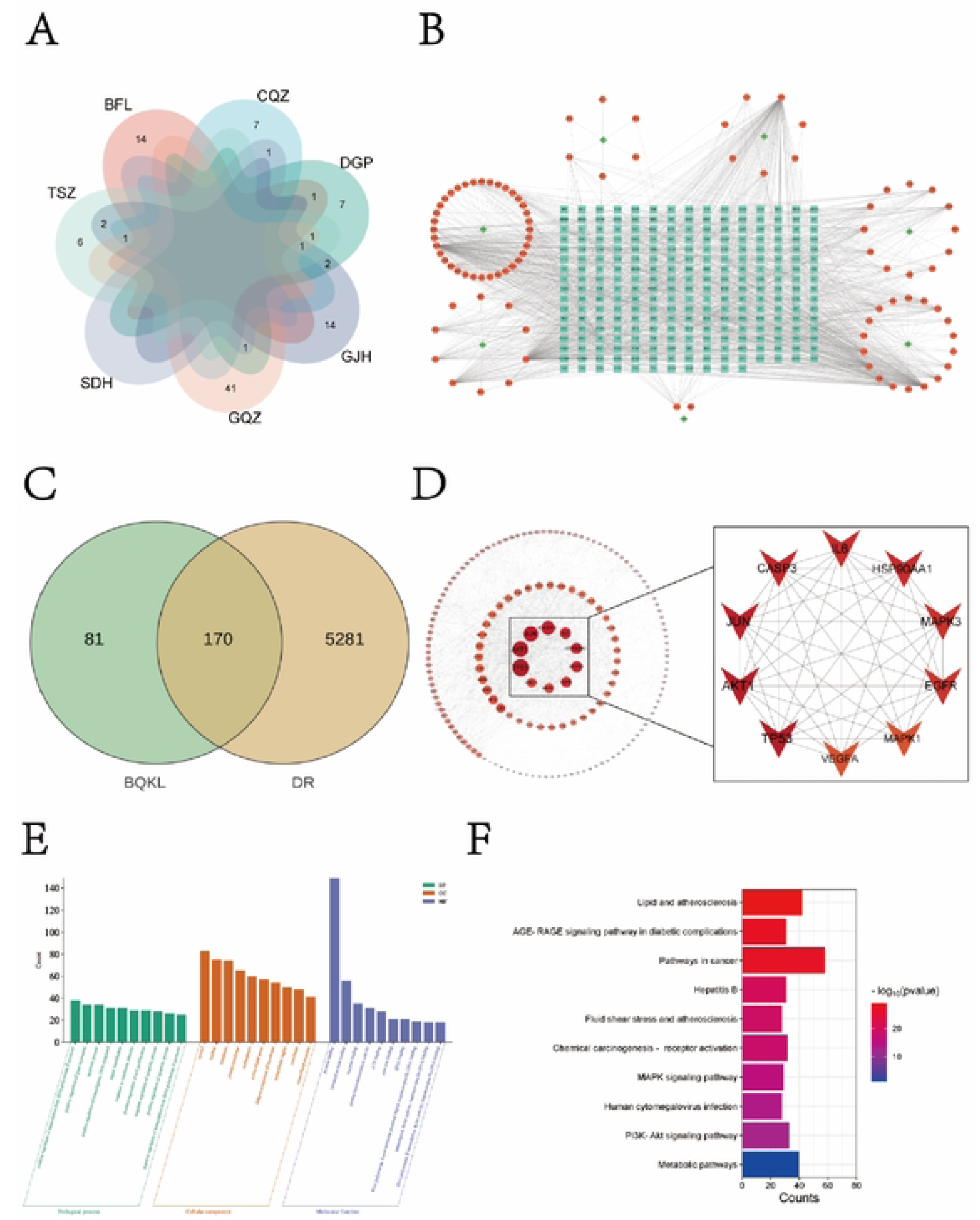
Prediction of network pharmacology. **(A)** Main Ingredients of BQKL. (B) “BQKL • active ingredient - common target networl(’ diagram. The rhombus represents the herbal medicine, the square hexagon is the active ingredient, and the square is the target site. (C) Venn diagram of BQKL and DR intersection targets. (D) PPI network diagram of BQKL therapeutic DR targets of action. **(E)** GO enrichment analysis results. (F) KEGG GOenrichment analysis results.

### 3.2. Potential targets for DR treatment with BQKL

By searching the above disease databases, a total of 5451 relevant targets were obtained after merging and de-emphasizing. Using the software to identify the intersection of DR targets with the compound targets of BQKL, a total of 170 potential targets for the treatment of DR were obtained (Fig 2C).

### 3.3. Construction of the PPI network

The results of String database analysis were further analyzed and visualized by Cytoscape software to establish the PPI network. From the figure, it can be found that TP53, AKT1, JUN, CASPASE3, MAPK3 and MAPK1 are closely related to other targets (Fig 2D).

### 3.4. Target enrichment analysis

Among them, a total of 727 pathways were obtained by GO analysis for biological processes, which mainly involved apoptotic processes, inflammatory responses, responses to hypoxia, etc. 93 pathways for molecular functions, which mainly involved cytoplasm, nucleus, cytoplasm, etc. and 153 pathways for cellular compositions, which mainly included protein-binding, binding of the same proteins, and enzyme-binding (Fig 2E). A total of 176 pathways were obtained by KEGG analysis, which mainly involved the cancer pathway, the AGE-RAGE signaling pathway in diabetic complications, the metabolic pathway, the MAPK signaling pathway, the PI3K-AKT signaling pathway, and other key signaling pathways (Fig 2F).

### 3.5. Molecular docking

The lower the free energy of binding of molecules and targets to each other, the more stable the binding between ligand and receptor proteins. Molecular docking analysis showed that quercetin, kaempferol, β-sitosterol, lignanserin and stigmasterol all formed stable docking with the core proteins TP53, AKT1, JUN, CASPASE3, MAPK3 and MAPK1, and their binding energies were well lower than -5.0 kCal•mol^-1^. The visualization of molecular docking results was completed by Pymol software (Fig 3).

**Fig 3.**
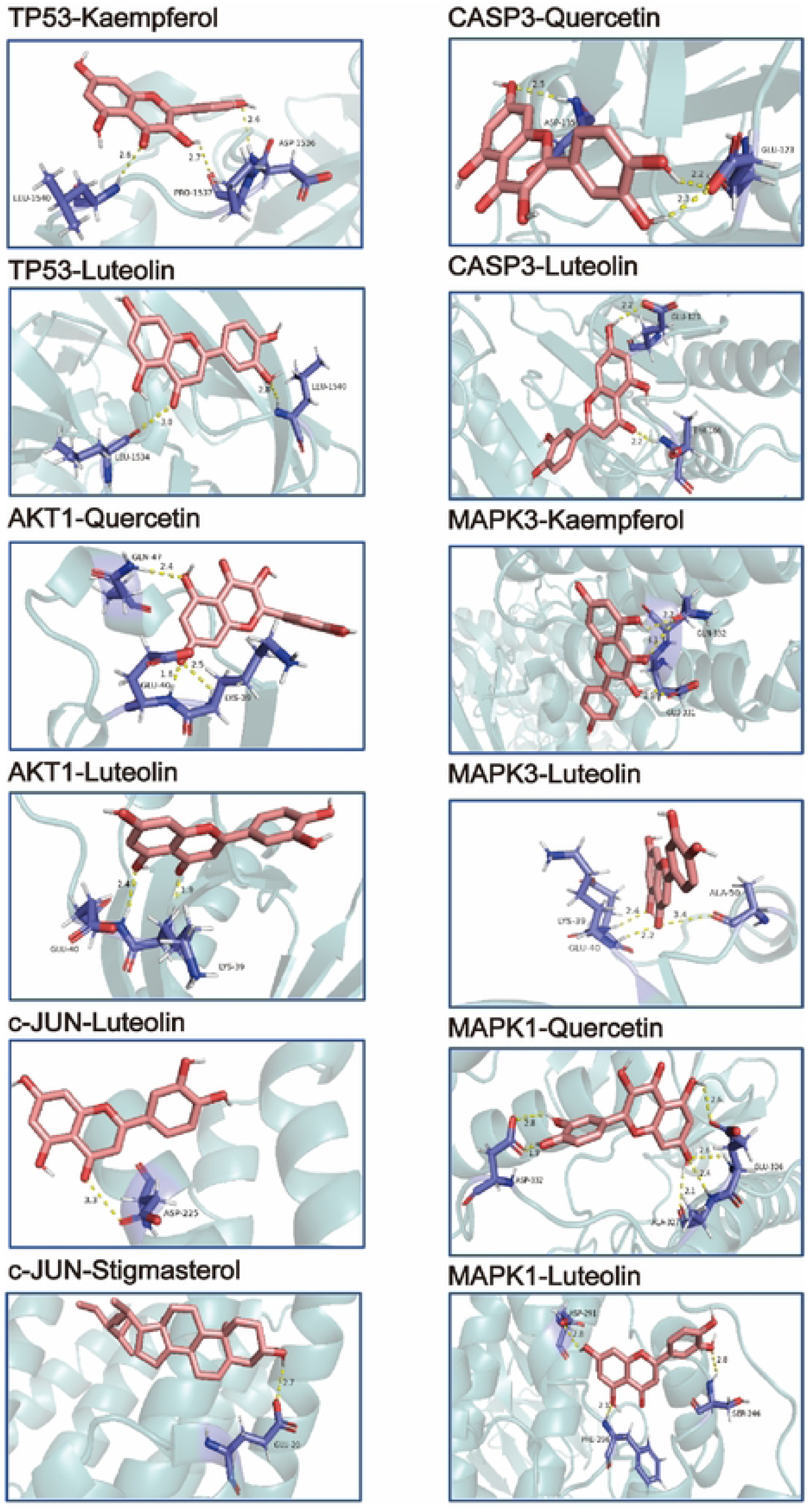
Molecular Docking.

**Table 2.** Molecular docking binding energy.

| Molecule Name | Binding energy (kcal/mol) |  |  |  |  |  |
| --- | --- | --- | --- | --- | --- | --- |
|  | TP53 | AKT1 | JUN | CASPASE3 | MAPK3 | MAPK1 |
| Quercetin | -7.3 | -6.4 | -6.4 | -7.9 | -7.9 | -8.4 |
| Kaempferol | -7.1 | -6 | -6.3 | -7.9 | -8.1 | -7.9 |
| Beta-sitosterol | -7.6 | -7.2 | -6.1 | -7.3 | -8.7 | -7.8 |
| Luteolin | -7.2 | -6.5 | -6.5 | -8.6 | -8.2 | -8.3 |
| Stigmasterol | -7 | -7.7 | -6.9 | -7.6 | -7 | -8.1 |

### 3.6. Effects of BQKL on general state, body mass and blood glucose in DR Rats

The rats in the normal control group exhibited good mental state, responsive behavior, soft hair, normal intake of diet and water, regular urine volume, and dry bedding throughout the experimental process. The rats in the model group displayed lethargy, disheveled and yellowish hair prone to depilation, significantly increased intake of diet and water as well as urine volume, along with moist bedding material. The rats in the treatment group showed some improvement compared to the model group.

Before modeling, there were no significant differences in body weight and blood glucose levels among all groups. After modeling, the control group exhibited a steady increase in body weight, while the model group showed no obvious changes and experienced varying degrees of decrease over time *(P <0.001)*. Additionally, fasting blood glucose levels increased significantly in the model group *(P < 0.001)*. At 8, 12, and 16 weeks, the model group’s body weight began to decrease significantly *(P < 0.001)*, while their fasting blood glucose levels remained significantly higher than those of the control group *(P < 0.001)*. Compared to the model group, both BQKL and CaD groups demonstrated significant increases in body weight and significant decreases in fasting blood glucose levels (*P < 0.05*). (Fig 4A and 4B).

**Fig 4.**
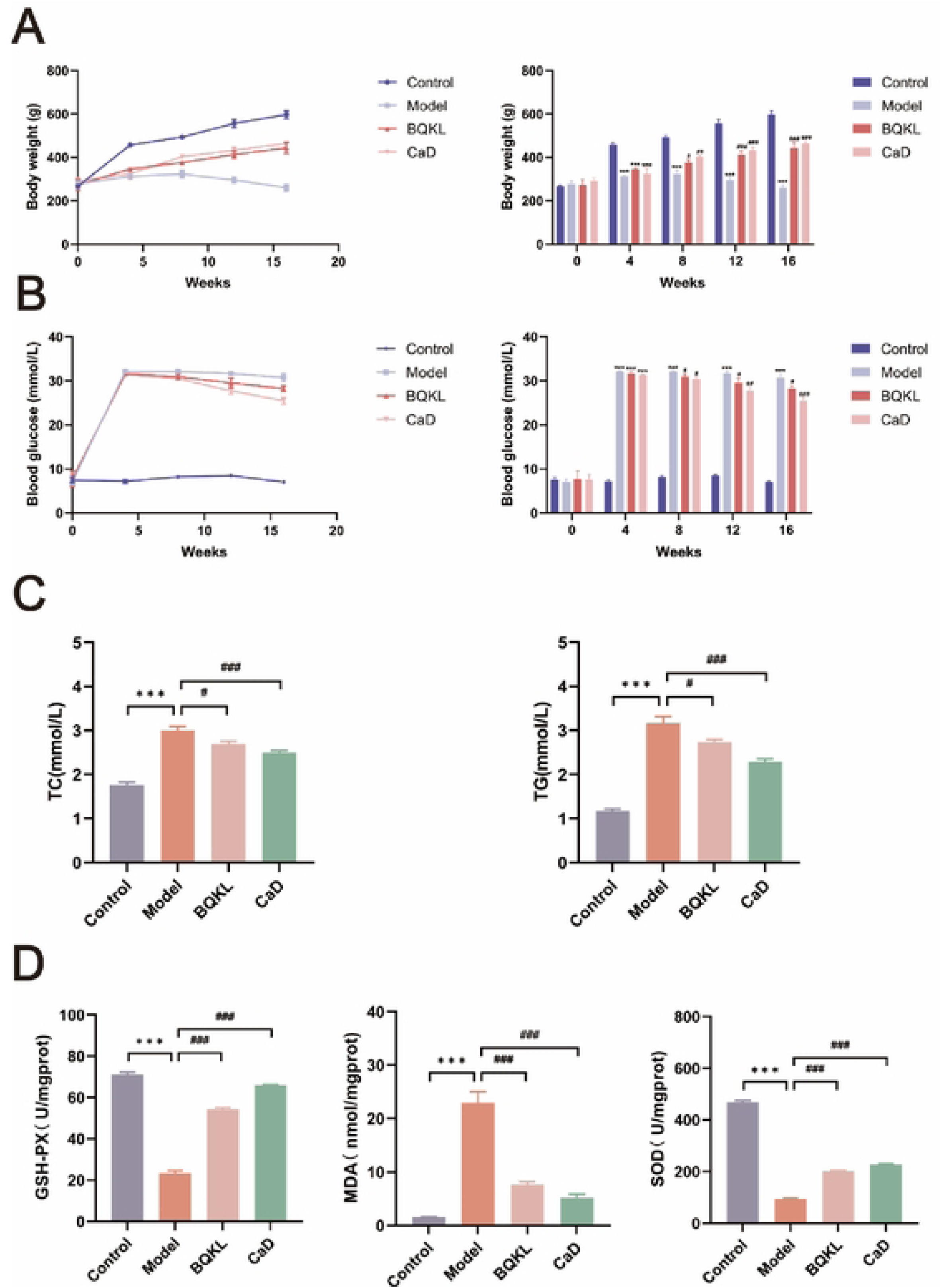
Metabolic parameters in DR rats. (A) Body weight at different time points. (B) Blood glucose at different time points. (C) Effect of BQKL on TC and TG levels in serum of DR rats. (D) Effect of BQKL on SOD, MDA and GPx contents in the retina of DR rats. ("p < 0.05: vs. NC group; # p < 0.05: vs. DR group)

### 3.7. Effects of BQKL on serum levels of TC, TG, SOD, MDA, and GPx in DR rats

Compared with the control group, the serum levels of TC, TG, and MDA were significantly higher, and the levels of SOD and GPx were significantly lower in the DR group rats (*P < 0.001*). Compared with the model group, the serum levels of TC, TG, and MDA were significantly lower, and the levels of SOD and GPx were significantly higher in the BQKL and CaD groups of rats (*P < 0.05*) (Fig 4C and 4D).

### 3.8. Histopathologic and morphological manifestations of the retina in each group of rats

In the control group, the structure of each retinal layer was clear and intact, and the cells in each layer were tightly arranged with normal morphology. There were fewer ganglion cells, loose cells in all layers, an unorganized arrangement, and vacuolar changes in the ganglion cell layer in the model group compared to the normal group (*P < 0.001*). Compared with the model group, the thickness of the retina of rats in the BQKL Group and the CaD Group was increased, the structure of each layer was neatly arranged, the hierarchy was clear, and the number of ganglion cells was increased (*P < 0.05*) (Fig 5).

**Fig 5.**
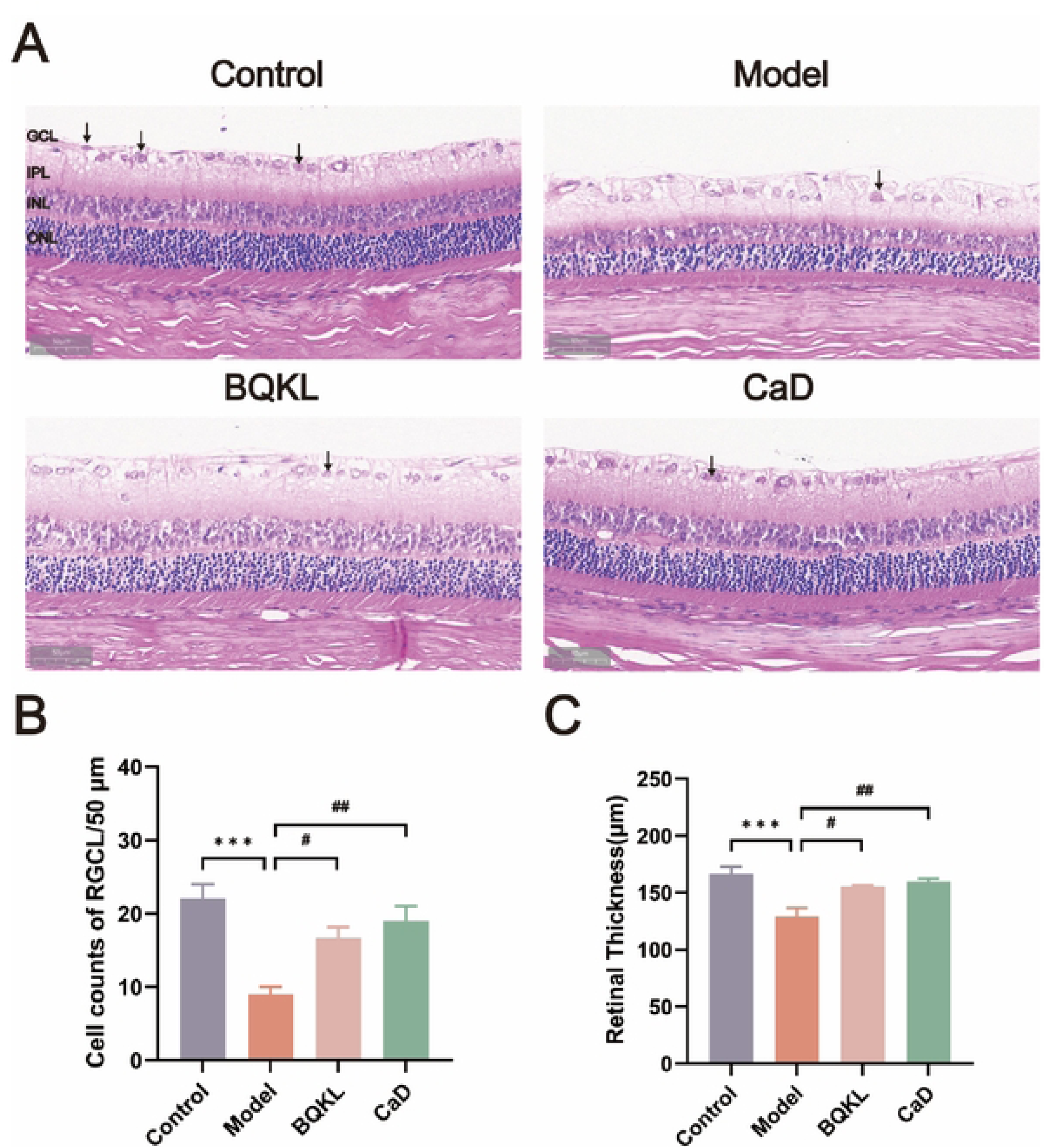
Effects of BQKL on retinal morphology in DR rats. (A) HE staining of the retina of rats in each group (•400-fold). (8) Count of retinal ganglion cells in each group. (C) Comparison of retinal thickness in each group. (*p < 0.05: vs. NC group; # p < 0.05: vs. DR group)

### 3.9. Immunohistochemical staining of CASPASE3, c-JUN, MAPK1, and MAPK3 in the retinal tissues of rats in each group

Compared to the control group, the model group exhibited significantly higher positive expression of rat CASPASE3 and c-JUN in retinal tissue (*P < 0.01*), while the positive expression of MAPK1 and MAPK3 was significantly lower (*P < 0.01*). In comparison to the model group, both BQKL and CaD groups showed decreased positive expression of rat CASPASE3 and c-JUN in retinal tissues (*P < 0.01*), along with elevated positive expression of MAPK1 and MAPK3 (*P < 0.05*) (Fig 6).

**Fig 6.**
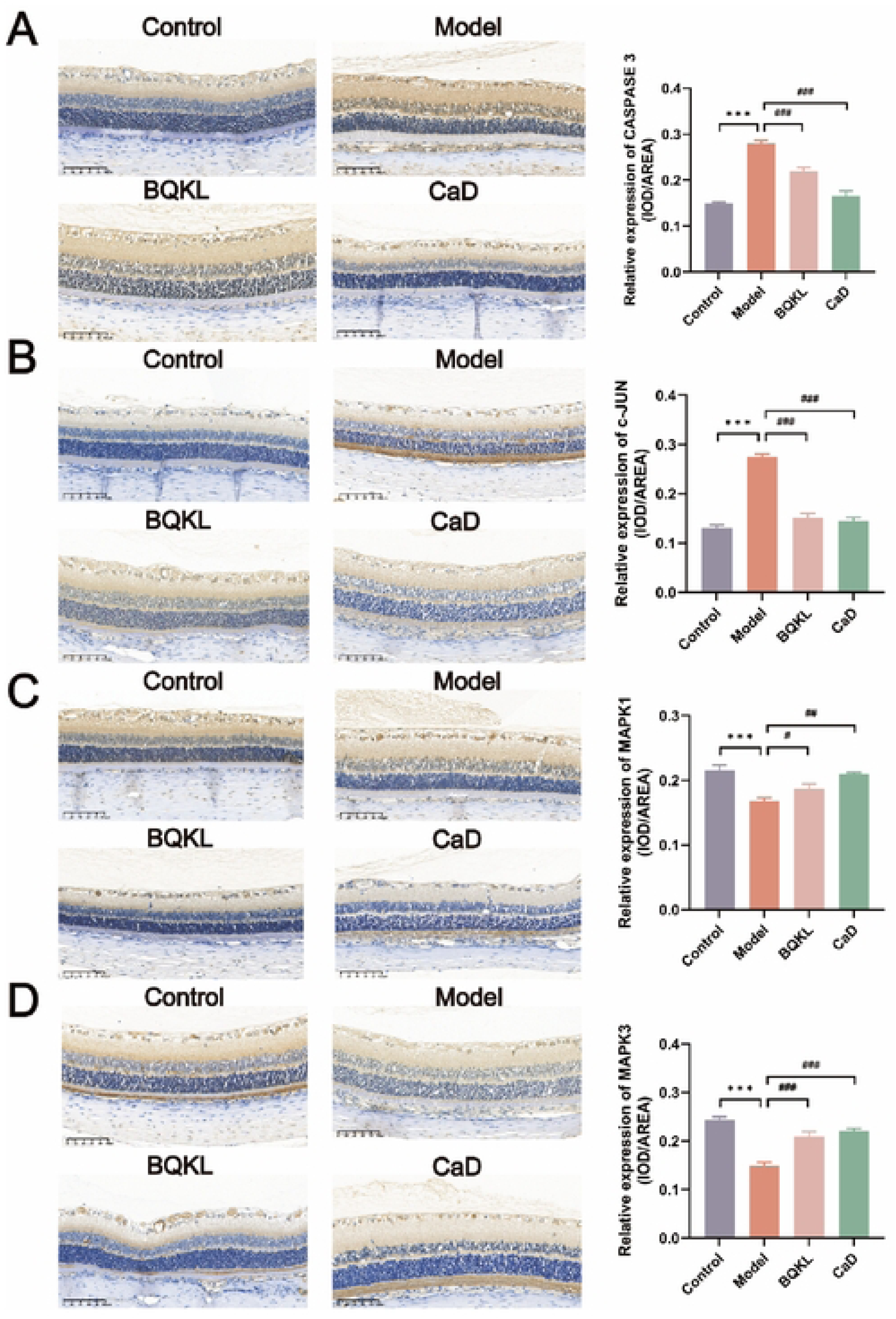
Immunohistochemical results of different groups of rat retinal tissue. (A) Immunohistochemical detection of CASPASE3 positive expression (•200-fold). (B) immunohistochemical detection of c-JUN positive expression (•200-fold). (C) Immunohistochemical detection of MAPK1 positive expression (•200-fold). (D) Immunohistochemical detection of MAPK3 positive expression (•200-fold). ("p < 0.05: vs. NC group; # p < 0.05: vs. DR group)

### 3.10. *CASPASE3*, *AKT1*, *c-JUN*, *TP53*, *MAPK1*, *MAPK3* mRNA expression in the rat retina of each group

The results showed that Caspase3, c-JUN, and TP53 mRNA in the retinal tissues of rats in the model group were significantly elevated compared with the control group (P < 0.05), and AKT1, MAPK1, and MAPK3 mRNA were significantly decreased (P < 0.05). The mRNA expressions of Caspase3, c-JUN, and TP53 in the BQKL group and CaD group were significantly lower than those in the model group (P < 0.05), while the mRNA expressions of AKT1, MAPK1, and MAPK3 were significantly higher (P < 0.05) (Fig 7A).

**Fig 7.**
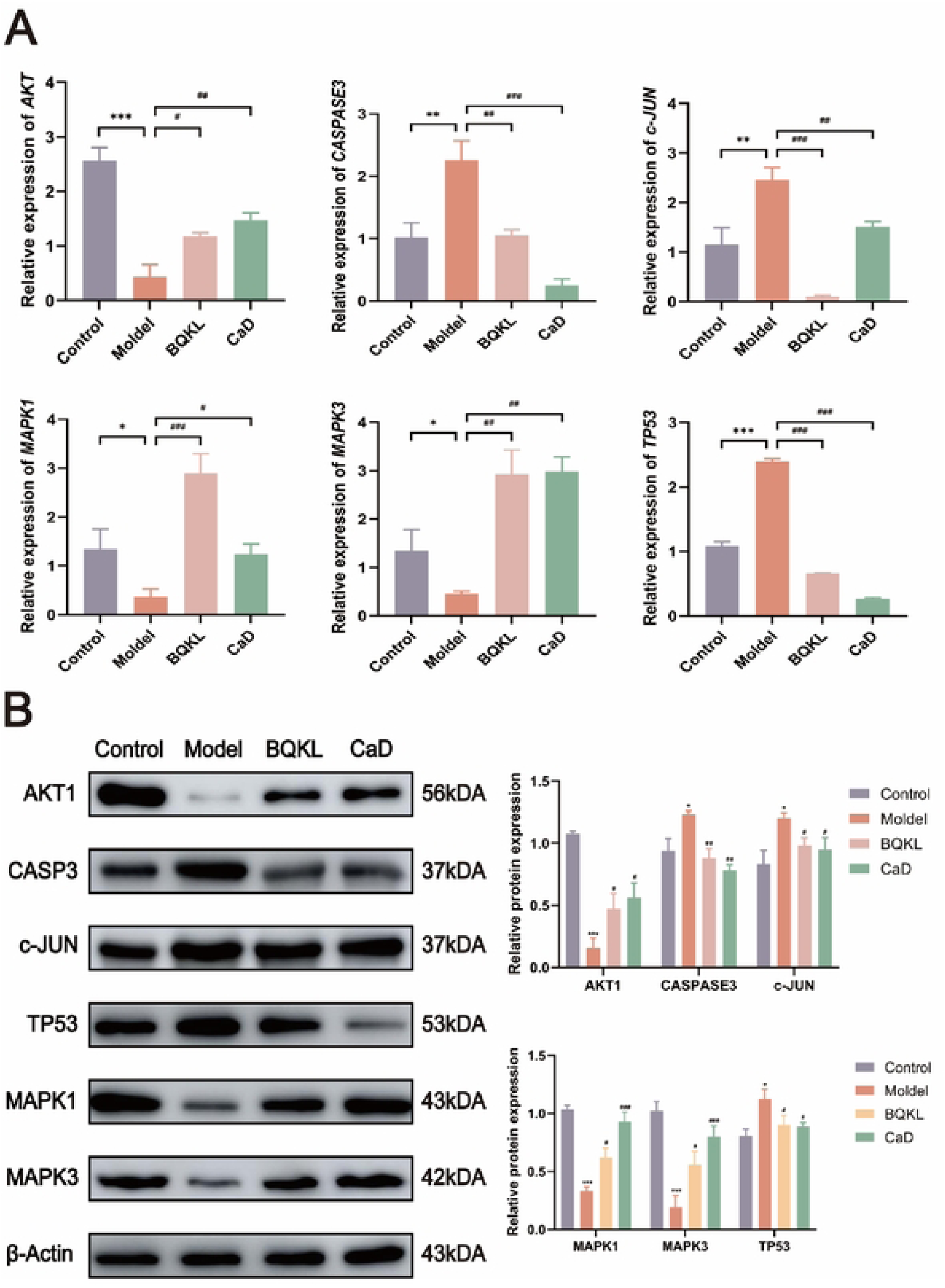
A molecular biological perspective on the role of BQKL in the treatment of DR. (A) The mRNA expression levels of *CASPASE3, AKT1, c.JUN, TP53, MAPK1,* and *MAPK3* were determined byqRT-PCR. (B) The protein expression levels of CASPASE3, AKT1, c-JUN, TP53, MAPK1, and MAPK3 were determined by Western blot. (*p < 0.05: vs. NC group; # p < 0.05: vs. DR group)

### 3.11. Protein expression levels of CASPASE3, AKT1, c-JUN, TP53, MAPK1, and MAPK3 in the retina of rats in each group

In the model group of rats, the levels of CASPASE3, c-JUN, and TP53 proteins were significantly higher (*P < 0.05*) than in the control group. On the other hand, the levels of AKT1, MAPK1, and MAPK3 proteins were significantly lower (*P < 0.05*). In contrast with the model group, the expression levels of CASPASE3, c-JUN, and TP53 proteins in the retinal tissues of rats in the BQKL group and the CaD group were significantly lower (*P < 0.05*), and the expression levels of AKT1, MAPK1, and MAPK3 proteins were significantly higher (*P < 0.05*) (Fig 7B).

## 4. Discussion

The network pharmacology technique was utilized to identify 97 active ingredients from 7 Chinese herbal medicines of BQKL. The prominent ingredients included quercetin, kaempferol, β-sitosterol, lignanserin, and stigmasterol. In recent years, certain Traditional Chinese Medicine have demonstrated significant hypoglycemic effects on diabetic patients and animal models, garnering considerable attention. Due to their naturalness and safety, these medicines are widely employed in clinical practice. Research has shown that the above ingredients have a variety of effects, such as lowering blood pressure, lowering blood lipids, lowering blood sugar, and having antioxidant and anti-inflammatory effects [30-32]. Quercetin was found to improve diabetes by enhancing lipid metabolism through antioxidant, anti-inflammatory, and activation of the FXR1/TGR5 signaling pathways [33]. Animal experiments have verified that kaempferol possesses excellent antioxidant properties and can significantly enhance the antioxidant status while reducing hyperglycemia in STZ-induced diabetic rats, thereby reducing the risk of diabetic complications [34]. β-Sitosterol controls hyperglycemia by activating the insulin receptor and glucose transporter protein 4 in adipocytes of type 2 diabetic rats and attenuates the inflammatory response in diabetic rats by down-regulating IKKβ/NF-κB and JNK signaling pathways [35]. By investigating the antidiabetic potential of lignocaine in STZ-induced diabetic rats following oral administration, it was discovered that lignocaine could reverse oxidative stress and inflammation, confirming its antioxidant and anti-inflammatory effects [36]. One study suggested that stigmasterol may improve insulin resistance by enhancing the expression and translocation of GLUT4, thus possibly exerting a hypoglycemic effect. Therefore, we can hypothesize that BQKL may delay the progression of DR through hypoglycemic, anti-inflammatory, and antioxidant mechanisms.

Adoption of databases on various diseases, a total of 5451 drug-related targets were obtained after merging and de-emphasizing. Through the construction of the PPI network, 170 potential therapeutic targets closely associated with the treatment of DR with BQKL were identified. The key core targets include TP53, AKT1, JUN, CASPASE3, MAPK3, and MAPK1. A relevant study discovered that excessive production of reactive oxygen species under hyperglycemic conditions can trigger inflammatory responses, mitochondrial dysfunction, and apoptosis, thereby influencing the onset and progression of DR [37]. TP53 is an anti-apoptotic transcription factor. It has been found that the activation of oxidative stress in retinal tissues is an important mechanism that promotes the development of fundopathy in diabetic patients, and that dichloroacetate can prevent DR through its anti-apoptotic, antioxidant, and antidiabetic properties [38]. A mouse model of STZ-induced DR demonstrated that AKT signaling in the retina may play a role in maintaining retinal homeostasis. Increasing Akt1 activity in the retina was found to reduce diabetes-induced molecular changes in the retina. In vitro and in vivo experiments were conducted to investigate the protective effect of hypericin against retinal damage caused by diabetic hyperglycemia [39]. The results showed that hypericin treatment significantly reduced the expression of apoptotic proteins CASPASE3, CASPASE9, and Bax, while increasing the expression of the anti-apoptotic protein Bcl-2 in the vascular endothelial cells of rats with diabetics [40]. Another study revealed that increasing EDEM1 blocked the IRE1/JNK/c-JUN signaling pathway, leading to elevated insulin mRNA levels. This improvement in insulin secretion normalized blood sugar and glucose tolerance levels in diabetic rats. Furthermore, human retinal endothelial cells treated with high glucose to mimic DR showed that overexpression of SPRED2 down-regulated the expression of p-ERK1/2 in HG-treated human retinal endothelial cells. This inhibition of inflammation and apoptosis not only alleviated the symptoms of DR but also protected the blood retinal barrier (BRB) in DR, thereby maintaining BRB homeostasis [41].

The GO enrichment analysis showed that the target genes for treating DR with BQKL were involved in biological processes like apoptosis, inflammation, response to hypoxia, and other things. This suggests that BQKL may have a lot of potential to help treat DR in many different ways. KEGG signaling pathway analysis showed that the therapeutic effect of BQKL on DR is mainly mediated through antioxidant, anti-inflammatory, and anti-apoptotic signaling pathways, mainly through the PI3K-AKT signaling pathway, the MAPK signaling pathway, and the AGE-RAGE signaling pathway. The AKT signaling pathway plays a crucial role in the development of DR in various retinal cells. A combination of in vitro and in vivo experiments has shown that it is possible to protect the integrity of the BRB by inhibiting the PI3K/Akt/Stat3/NF-κB signaling pathway, suppressing the overactivity of microglial cells, thereby decreasing the release of inflammation-inducing factors, and protecting the pericytes and endothelial cells [42]. Chronic hyperglycemia leads to non-enzymatic glycosylation of proteins and the formation of advanced glycosylation end-products (AGEs), which induce inflammation and immunosuppression by binding to AGE receptors, generating pro-inflammatory cytokines, reactive oxygen species, and reactive nitrogen intermediates, thereby affecting innate and adaptive immune responses [43]. The MAPK/ERK pathway is closely associated with cell proliferation, differentiation, and apoptosis [44]. Studies have shown that high glucose activates MAPK/ERK signaling in human umbilical vein endothelial cells, and human umbilical cord mesenchymal stem cells improve blood glucose levels in diabetic rats by reversing the abnormal phosphorylation of ERK in the MAPK/ERK signaling pathway, thereby protecting the vascular endothelium from diabetic damage [45].

We found that BQKL could effectively reduce blood glucose, blood lipids, and oxidative damage in diabetic rats, and HE staining results proved that BQKL could delay the development of DR, improve the morphology of the retina, and restore the thickness of the retina and the number of RGCs. Immunohistochemistry, Western blot, and qRT-PCR were used to examine the expression levels of CASPASE3, AKT1, c-JUN, TP53, MAPK1, and MAPK3 in retinal tissues, and the results showed that BQKL significantly up-regulated the expression of AKT1, MAPK1, and MAPK3 and down-regulated the expression of CASPASE3 in retinal tissues, c-JUN, and TP53 expression in a dose-dependent manner. This is further corroborated by the results of network pharmacology analysis, suggesting that BQKL can exert therapeutic effects by targeting CASPASE3, AKT1, c-JUN, TP53, MAPK1, and MAPK3, which are pharmacological targets.

## 5. Conclusion

In summary, BQKL has the advantages of multi-pathways and multi-targets in the treatment of DR. BQKL mainly improves oxidative stress, apoptosis, and inflammation due to hyperglycemia-related stress by regulating key targets such as CASPASE3, AKT1, c-JUN, TP53, MAPK1, MAPK3, and thus slows down the occurrence and development of DR (Fig 8). This paper further elucidates the molecular biological mechanism of BQKL for the treatment of DR through network pharmacological methods and animal experimental validation, which lays a solid theoretical foundation for the clinical treatment of DR. However, this study is only a preliminary investigation of the mechanism of action of BQKL in DR based on network pharmacology, and further experimental validation is still needed. In order to more effectively investigate the overall gene expression regulation pattern in organisms and to explore its connection with disease occurrence and development, we will analyse the complex genome more comprehensively through proteomics, metabolomics with high-throughput analysis technology, molecular interactions, etc., so that we hope to bring new ways for the diagnosis and treatment of DR.

**Fig 8.**
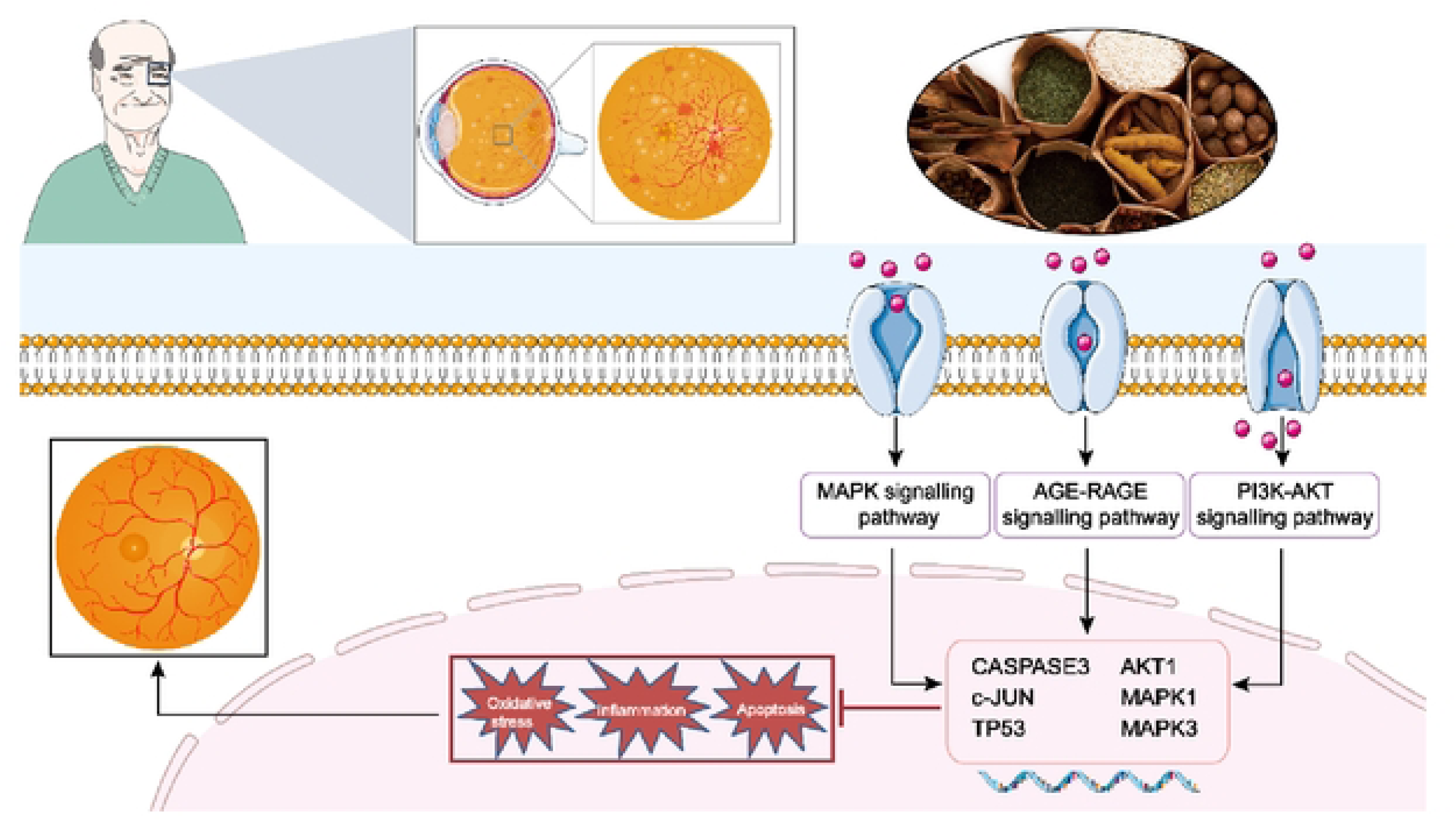
Graphical Abstract.

## Abbreviation

DR: Diabetic retinopathy
BQKL: Buqing Granule
GO: Gene Ontology
KEGG: Kyoto Encyclopaedia of Genes and Genomes
SPF: specific-pathogen-free
SD: Sprague Dawley
TSZ: *Cuscuta chinensis* Lam. (Tusizi)
GQZ: *Lycium barbarum* L. (Gouqizi)
DGP: *Lycium chinense* Mill. (Digupi)
CQZ: *Plantago asiatica* L. (Cheqianzi)
GJH: *Chrysanthemum morifolium* Ramat. (Ganjuhua)
SDH: *Rehmannia glutinosa* Libosch. (Shudihuang)
BFL: *Poria cocos* (Schw.) Wolf (Baifuling)
CaD: calcium dobesilate
STZ: streptozotocin
TC: cholesterol
TG: triglyceride
SOD: superoxide dismutase
MDA: malondialdehyde
GPx: glutathione peroxidase
HE: Hematoxylin-Eosin
RGC: retinal ganglion cells
BRB: blood retinal barrier
AGE: advanced glycosylation end-product

## References

1. Liu Y, Wu N. Progress of Nanotechnology in Diabetic Retinopathy Treatment. International journal of nanomedicine. 2021; 16: 1391–403.

2. Wong TY, Cheung CM, Larsen M, Sharma S, Simó R. Diabetic retinopathy. Nature reviews Disease primers. 2016; 2: 16012.

3. Wang W, Lo ACY. Diabetic Retinopathy: Pathophysiology and Treatments. International journal of molecular sciences. 2018; 19(6).

4. Cao H. Overseas Returned Chinese Medicine Ancient Books Collection. 12, Yangshi Jiacang Fang. Chinese Ancient Books Publishing Co; 2005.

5. Yu Y, Feng Z, Zhao X. Niu Y. Clinical observation on 35 cases of immature age-related cataract treated with Buqing soup. Journal of Ningxia Medical University. 2011, 33(5): 496–497.

6. Deng Z, Niu Y, Wang R, Fan Q, Lang S. Experimental Study on Protection and Treatment of Galactose Cataract by Buqing Granules. Chinese Journal of Experimental Traditional Medical Formulae. 2013, 19 (13): 205–208.

7. LI W, Niu Y, Nan Y, Yuan L, Ma H. Effects of Buqing Granules on mitochondrial autophagy Bnip3/Nix signaling pathway in high glucose induced human lens epithelium cells. China Journal of Traditional Chinese Medicine and Pharmacy. 2018,33(07): 3056-3060.

8. Yang J, Zhou H, Niu Y. Analysis of the chemical constituents of Buqing Soup by HPLC-IT-TOF method. Journal of Ningxia Medical University. 2018,40(2):235-238.

9. Lu Y, Pan X, Yuan L, Niu Y, Nan Y. Effect of Buqing Granule on Early and Middle Stage Diabetic Cataract in db/db Mice. Genomics and Applied Biology. 2021,40(Z3):3330-3336.

10. Fu H, Li W, Weng Z, Huang Z, Liu J, Mao Q, et al. Water extract of cacumen platycladi promotes hair growth through the Akt/GSK3β/β-catenin signaling pathway. Frontiers in pharmacology. 2023; 14:1038039.

11. Gan X, Shu Z, Wang X, Yan D, Li J, Ofaim S, et al. Network medicine framework reveals generic herb-symptom effectiveness of traditional Chinese medicine. Science advances. 2023;9(43): eadh0215.

12. Ru J, Li P, Wang J, Zhou W, Li B, Huang C, et al. TCMSP: a database of systems pharmacology for drug discovery from herbal medicines. Journal of cheminformatics. 2014; 6:13.

13. Pan S, Hu B, Sun J, Yang Z, Yu W, He Z, et al. Identification of cross-talk pathways and ferroptosis-related genes in periodontitis and type 2 diabetes mellitus by bioinformatics analysis and experimental validation. Frontiers in immunology. 2022; 13:1015491.

14. Amberger JS, Bocchini CA, Schiettecatte F, Scott AF, Hamosh A. OMIM.org: Online Mendelian Inheritance in Man (OMIM®), an online catalog of human genes and genetic disorders. Nucleic acids research. 2015;43(Database issue): D789–98.

15. Zhou Y, Zhang Y, Zhao D, Yu X, Shen X, Zhou Y, et al. TTD: Therapeutic Target Database describing target druggability information. Nucleic acids research. 2023.

16. Mitra-Ghosh T, Callisto SP, Lamba JK, Remmel RP, Birnbaum AK, Barbarino JM, et al. PharmGKB summary: lamotrigine pathway, pharmacokinetics and pharmacodynamics. Pharmacogenetics and genomics. 2020;30(4):81–90.

17. Barrett T, Wilhite SE, Ledoux P, Evangelista C, Kim IF, Tomashevsky M, et al. NCBI GEO: archive for functional genomics data sets--update. Nucleic acids research. 2013;41(Database issue): D991-5.

18. Heberle H, Meirelles GV, da Silva FR, Telles GP, Minghim R. InteractiVenn: a web-based tool for the analysis of sets through Venn diagrams. BMC bioinformatics. 2015;16(1):169.

19. Szklarczyk D, Gable AL, Nastou KC, Lyon D, Kirsch R, Pyysalo S, et al. The STRING database in 2021: customizable protein-protein networks, and functional characterization of user-uploaded gene/measurement sets. Nucleic acids research. 2021;49(D1): D605–d12.

20. Zhang J, Zhou Y, Ma Z. Multi-target mechanism of Tripteryguim wilfordii Hook for treatment of ankylosing spondylitis based on network pharmacology and molecular docking. Annals of medicine. 2021;53(1):1090–8.

21. Kim S. Getting the most out of PubChem for virtual screening. Expert opinion on drug discovery. 2016;11(9):843–55.

22. Rubach P, Zajac S, Jastrzebski B, Sulkowska JI, Sułkowski P. Genus for biomolecules. Nucleic acids research. 2020;48(D1): D1129–d35.

23. Chen M, Lv H, Gan J, Ren J, Liu J. Tang Wang Ming Mu Granule Attenuates Diabetic Retinopathy in Type 2 Diabetes Rats. Frontiers in physiology. 2017; 8:1065.

24. Liang WJ, Yang HW, Liu HN, Qian W, Chen XL. HMGB1 upregulates NF-kB by inhibiting IKB-α and associates with diabetic retinopathy. Life sciences. 2020; 241:117146.

25. Pang B, Li M, Song J, Li QW, Wang J, Di S, et al. Luo Tong formula attenuates retinal inflammation in diabetic rats via inhibition of the p38MAPK/NF-κB pathway. Chinese medicine. 2020; 15:5.

26. Sadikan MZ, Nasir NAA, Iezhitsa I, Agarwal R. Antioxidant and anti-apoptotic effects of tocotrienol-rich fraction against streptozotocin-induced diabetic retinopathy in rats. Biomedicine & pharmacotherapy = Biomedecine & pharmacotherapie. 2022; 153:113533.

27. Xiang E, Han B, Zhang Q, Rao W, Wang Z, Chang C, et al. Human umbilical cord-derived mesenchymal stem cells prevent the progression of early diabetic nephropathy through inhibiting inflammation and fibrosis. Stem cell research & therapy. 2020;11(1):336.

28. Wang X, Pan J, Liu H, Zhang M, Liu D, Lu L, et al. AIM2 gene silencing attenuates diabetic cardiomyopathy in type 2 diabetic rat model. Life sciences. 2019; 221:249–58.

29. Sun HH, Chai XL, Li HL, Tian JY, Jiang KX, Song XZ, et al. Fufang Xueshuantong alleviates diabetic retinopathy by activating the PPAR signalling pathway and complement and coagulation cascades. Journal of ethnopharmacology. 2021; 265:113324.

30. Hosseini A, Razavi BM, Banach M, Hosseinzadeh H. Quercetin and metabolic syndrome: A review. Phytotherapy research: PTR. 2021;35(10):5352–64.

31. Babu S, Jayaraman S. An update on β-sitosterol: A potential herbal nutraceutical for diabetic management. Biomedicine & pharmacotherapy = Biomedecine & pharmacotherapie. 2020; 131:110702.

32. Yang Y, Chen Z, Zhao X, Xie H, Du L, Gao H, et al. Mechanisms of Kaempferol in the treatment of diabetes: A comprehensive and latest review. Frontiers in endocrinology. 2022; 13:990299.

33. Yang H, Yang T, Heng C, Zhou Y, Jiang Z, Qian X, et al. Quercetin improves nonalcoholic fatty liver by ameliorating inflammation, oxidative stress, and lipid metabolism in db/db mice. Phytotherapy research: PTR. 2019;33(12):3140–52.

34. Al-Numair KS, Chandramohan G, Veeramani C, Alsaif MA. Ameliorative effect of kaempferol, a flavonoid, on oxidative stress in streptozotocin-induced diabetic rats. Redox report: communications in free radical research. 2015;20(5):198–209.

35. Kahksha, Alam O, Al-Keridis LA, Khan J, Naaz S, Alam A, et al. Evaluation of Antidiabetic Effect of Luteolin in STZ Induced Diabetic Rats: Molecular Docking, Molecular Dynamics, In Vitro and In Vivo Studies. Journal of functional biomaterials. 2023;14(3).

36. Wang J, Huang M, Yang J, Ma X, Zheng S, Deng S, et al. Anti-diabetic activity of stigmasterol from soybean oil by targeting the GLUT4 glucose transporter. Food & nutrition research. 2017;61(1):1364117.

37. Kang Q, Yang C. Oxidative stress and diabetic retinopathy: Molecular mechanisms, pathogenetic role and therapeutic implications. Redox biology. 2020; 37:101799.

38. Kanan Y, Hackett SF, Taneja K, Khan M, Campochiaro PA. Oxidative stress-induced alterations in retinal glucose metabolism in Retinitis Pigmentosa. Free radical biology & medicine. 2022; 181:143–53.

39. Liu H, Stepicheva NA, Ghosh S, Shang P, Chowdhury O, Daley RA, et al. Reducing Akt2 in retinal pigment epithelial cells causes a compensatory increase in Akt1 and attenuates diabetic retinopathy. Nature communications. 2022;13(1):6045.

40. Wu W, Xie Z, Zhang Q, Ma Y, Bi X, Yang X, et al. Hyperoside Ameliorates Diabetic Retinopathy via Anti-Oxidation, Inhibiting Cell Damage and Apoptosis Induced by High Glucose. Frontiers in pharmacology. 2020; 11:797.

41. Flintoaca Alexandru PR, Chiritoiu GN, Lixandru D, Zurac S, Ionescu-Targoviste C, Petrescu SM. EDEM1 regulates the insulin mRNA level by inhibiting the endoplasmic reticulum stress-induced IRE1/JNK/c-Jun pathway. iScience. 2023;26(10):107956.

42. Tang L, Zhang C, Lu L, Tian H, Liu K, Luo D, et al. Melatonin Maintains Inner Blood-Retinal Barrier by Regulating Microglia via Inhibition of PI3K/Akt/Stat3/NF-κB Signaling Pathways in Experimental Diabetic Retinopathy. Frontiers in immunology. 2022; 13:831660.

43. Shen CY, Lu CH, Wu CH, Li KJ, Kuo YM, Hsieh SC, et al. The Development of Maillard Reaction, and Advanced Glycation End Product (AGE)-Receptor for AGE (RAGE) Signaling Inhibitors as Novel Therapeutic Strategies for Patients with AGE-Related Diseases. Molecules (Basel, Switzerland). 2020;25(23).

44. Fang JY, Richardson BC. The MAPK signalling pathways and colorectal cancer. The Lancet Oncology. 2005;6(5):322–7.

45. Liu Y, Chen J, Liang H, Cai Y, Li X, Yan L, et al. Human umbilical cord-derived mesenchymal stem cells not only ameliorate blood glucose but also protect vascular endothelium from diabetic damage through a paracrine mechanism mediated by MAPK/ERK signaling. Stem cell research & therapy. 2022;13(1):258.

